# 2’-Fucosyllactose Inhibits Human Norovirus Replication in Human Intestinal Enteroids

**DOI:** 10.1101/2024.05.30.596597

**Authors:** Ketki Patil, B. Vijayalakshmi Ayyar, Frederick H. Neill, Lars Bode, Mary K. Estes, Robert L. Atmar, Sasirekha Ramani

**Affiliations:** Department of Molecular Virology and Microbiology, Baylor College of Medicine, Houston, Texas; Department of Pediatrics, Larsson-Rosenquist Foundation Mother-Milk-Infant Center of Research Excellence (MOMI CORE), and the Human Milk Institute (HMI), University of California San Diego, La Jolla, CA; Department of Medicine, Baylor College of Medicine, Houston, Texas

**Keywords:** Norovirus, Human Milk Oligosaccharide, 2’-Fucosyllactose, Enteroids, Antiviral, Therapeutic

## Abstract

Human noroviruses (HuNoVs) are the leading cause of acute gastroenteritis worldwide. Currently, there are no targeted antivirals for the treatment of HuNoV infection. Histo-blood group antigens (HBGAs) on the intestinal epithelium are cellular attachment factors for HuNoVs; molecules that block the binding of HuNoVs to HBGAs thus have the potential to be developed as antivirals. Human milk oligosaccharides (HMOs) are glycans in human milk with structures analogous to HBGAs. HMOs have been shown to act as decoy receptors to prevent the attachment of multiple enteric pathogens to host cells. Previous X-ray crystallography studies have demonstrated the binding of HMO 2’-fucosyllactose (2’FL) in the same pocket as HBGAs for some HuNoV strains. We evaluated the effect of 2’FL on the replication of a globally dominant GII.4 Sydney [P16] HuNoV strain using human intestinal enteroids (HIEs) from adults and children. A significant reduction in GII.4 Sydney [P16] replication was seen in duodenal and jejunal HIEs from multiple adult donors, all segments of the small intestine from an adult organ donor and in two pediatric duodenal HIEs. However, 2’FL did not inhibit HuNoV replication in two infant jejunal HIEs that had significantly lower expression of α1-2-fucosylated glycans. 2’FL can be synthesized in large scale, and safety and tolerance have been assessed previously. Our data suggest that 2’FL has the potential to be developed as a therapeutic for HuNoV gastroenteritis.

**IMPORTANCE:** Human noroviruses infect the gastrointestinal tract and are a leading cause of acute gastroenteritis worldwide. Common symptoms of norovirus include diarrhea, vomiting and stomach cramps. Virus shedding and symptoms are prolonged and debilitating in immunocompromised patients. Currently, there are no approved vaccines or targeted antivirals for treating human norovirus infection. Human intestinal enteroids derived from intestinal stem cells allow the successful replication of norovirus in the laboratory and can be used as a physiologically relevant model system to evaluate antivirals. We discovered that 2’fucosyllactose (2’FL), an oligosaccharide naturally occurring in human milk, inhibits norovirus replication in HIEs from multiple donors and thus has the potential to be developed as a therapeutic for human norovirus. These findings have high translational potential since 2’FL from several manufacturers have GRAS (generally recognized as safe) status and can be synthesized on a large scale for immediate application.

## INTRODUCTION

Human noroviruses (HuNoVs) are a leading cause of acute gastroenteritis across all age groups (1). There are an estimated 677 million HuNoV infections worldwide and over 200,000 HuNoV-associated deaths each year, with the latter mainly reported in low- and middle-income countries (2, 3). HuNoV outbreaks have been reported in hospitals, long-term care facilities, cruise ships, planes and restaurants (4). Each year, HuNoV infections can result in more than $4 billion and $60 billion in direct health and societal care costs respectively (5). Currently, there are no targeted antivirals or licensed vaccines for HuNoVs.

Host cellular factors involved in virus attachment and entry are potential targets for antiviral development. Histo-blood group antigens (HBGAs) are cellular attachment factors for HuNoVs (6). These complex carbohydrates are present on red blood cells, mucosal epithelial cells, and biological fluids (7). Human milk contains a group of structurally diverse unconjugated glycans, with some structures analogous to HBGAs (8). These sugars, called human milk oligosaccharides (HMOs), comprise 5-15g/L of mature human milk and are the third most abundant component of human milk after lactose and lipids (9, 10). More than 150 HMO structures have been identified (11). In addition to serving as prebiotics for bacteria in the infant gut, other functions of HMOs include modulating epithelial and immune cell responses and acting as decoy receptors to reduce the attachment of pathogenic microbes to cell surface receptors (12). As such, HMOs have been shown to prevent pathogen adhesion to host epithelia for multiple enteric bacteria such as *Campylobacter jejuni, Clostridioides difficile* and *Escherichia coli* O157 as well as viruses such as rotavirus, coxsackievirus class A type 9 and SARS-CoV-2 (13–18).

Previous X-ray crystallography studies with three HuNoV genotypes (GI.1, GII.10 and GII.17) have shown that 2-fucosyllactose (2’FL), an α-1,2-fucosylated HMO, binds to the protruding domain of the HuNoV capsid protein VP1 in a similar pocket as HBGAs (19–21). 2’FL has also been found to block the binding of HuNoV virus-like particles (VLPs) to porcine gastric mucin (PGM) and saliva that contains HBGAs (19, 21). These data suggest that 2’FL can potentially act as a decoy receptor for HuNoVs. We previously standardized a pipeline to evaluate antivirals against HuNoVs in human intestinal enteroids (HIEs) (23). In the present study, we used this pipeline to evaluate the effect of 2’FL on the replication of GII.4 Sydney [P16] HuNoV and demonstrate significant reduction in HIEs from multiple donors and intestinal segments.

## RESULTS

### 2’FL SIGNIFICANTLY REDUCES GII.4 VLP BINDING TO PGM

We first carried out dose-response assays using different concentrations of 2’FL (1.25 mg/ml, 2.5 mg/ml, 5 mg/ml, 10 mg/ml and 20 mg/ml) to determine if the HMO used in the present study can reduce the binding of GII.4 Sydney 2012 VLPs to PGM. There was a dose-dependent reduction in VLP binding to PGM, with a significant reduction at 20 mg/ml 2’FL (Figure 1), suggesting that 2’FL can act as a decoy to block HuNoV replication.

**Figure 1:**
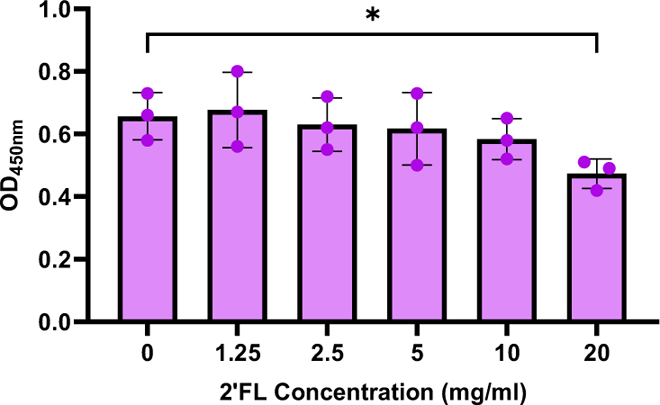
20 mg/ml 2’FL significantly reduces GII.4 Sydney 2012 VLP binding to PGM. Dose-response studies testing 1.25 mg/ml, 2.5 mg/ml, 5 mg/ml, 10 mg/ml and 20 mg/ml of 2’FL with 2.5 ug/ml VLPs. All comparisons were made to the condition where 2’FL was not present (0 mg/ml). Data represented are from n=3 independent experiments with averages from 3 technical replicates per experiment. The P-values were calculated using Student’s t-test. *P ≤ 0.05.

### HUMAN NOROVIRUS TISSUE CULTURE INFECTIOUS DOSE DIFFERS PER HIE LINE

We demonstrated previously that the number of genome equivalents (GE) per 50% tissue culture infectious dose (TCID_50_) of HuNoV strains differ in each HIE line (23). Since we planned to examine the effect of 2’FL in inhibiting HuNoV replication using HIE lines from different ages and intestinal segments, we determined the GE/TCID_50_ of the GII.4 Sydney [P16] HuNoV isolate in each line to standardize the amount of virus used across HIE lines. The average GE/TCID_50_ from two independent experiments are reported in Table 1, with adult duodenal HIEs requiring the highest number of GE/TCID_50_. For the duodenum and jejunum where HIEs from adults and children were available, the GE/TCID_50_ was lower in HIE lines from children. For HIEs derived from different intestinal segments of the same donor, the highest GE/TCID_50_ was seen in the duodenal HIE D2004 while the ileal line I2004 had lower GE/TCID_50_ values, almost similar to that of infant jejunal lines (J1005 and J1006). Taken together, these data indicate segment- and age-specific differences in GE/TCID_50_ and the need to standardize inoculum used in infectivity assays to allow for comparisons between HIE lines.

**Table 1:** Summary of genome equivalents (GE) per 50% tissue culture infectious dose (TCID_50_). List of HIE lines, segment of origin and age of donors are shown. GE/TCID_50_ values are shown as log_10_ values ± standard deviation (SD) from n=2 independent experiments.

| HIE | Segment | Age | GE/ TCID<sub>50</sub> ± SD |
| --- | --- | --- | --- |
| D109 | Duodenal | 44 years | 4.35 ± 0.28 |
| D2004 | Duodenal | 25 years | 4.54 ± 0.17 |
| J2 | Jejunal | 52 years | 4.05 ± 0.00 |
| J11 | Jejunal | 52 years | 4.06 ± 0.15 |
| J2004 | Jejunal | 25 years | 4.30 ± 0.03 |
| I2004 | Ileal | 25 years | 3.76 ± 0.06 |
| 4D | Duodenal | 2 years | 4.23 ± 0.23 |
| 8D | Duodenal | 5 years | 3.84 ± 0.21 |
| J1005 | Jejunal | 10 weeks | 3.78 ± 0.25 |
| J1006 | Jejunal | 12 weeks | 3.63 ± 0.06 |

### 2’FL SIGNIFICANTLY REDUCES GII.4 HUMAN NOROVIRUS REPLICATION IN ADULT DUODENAL HIE LINES

We first carried out dose-response assays to determine if 2’FL inhibited GII.4 Sydney [P16] HuNoV binding and replication in HIEs. Although significant inhibition of VLP binding to PGM was seen only with 20 mg/ml of 2’FL, we tested two additional concentrations (5 mg/ml and 10 mg/ml) to determine if lower doses could be effective in infectivity studies. HIE lines were infected with 100 TCID_50_ of GII.4 Sydney [P16] HuNoV based on their respective GE/TCID_50_ (Table 1). In the absence of 2’FL, GII.4 Sydney [P16] HuNoV showed ∼1log_10_ increase in GE/well at 24 hours post infection (hpi) compared to 1 hpi for D109 (Figure 2A) and D2004 (Figure 2B) HIE lines. Similar to the VLP studies, only 20 mg/ml of 2’FL significantly inhibited GII.4 Sydney [P16] HuNoV replication as measured at 24 hpi. In evaluating the effect of 2’FL on GII.4 Sydney [P16] HuNoV binding at 1 hpi, 20 mg/ml 2’FL significantly reduced binding in D2004 but not D109 HIE. None of the 2’FL concentrations tested were cytotoxic to HIEs as measured by the lactase dehydrogenase assay.

**Figure 2:**
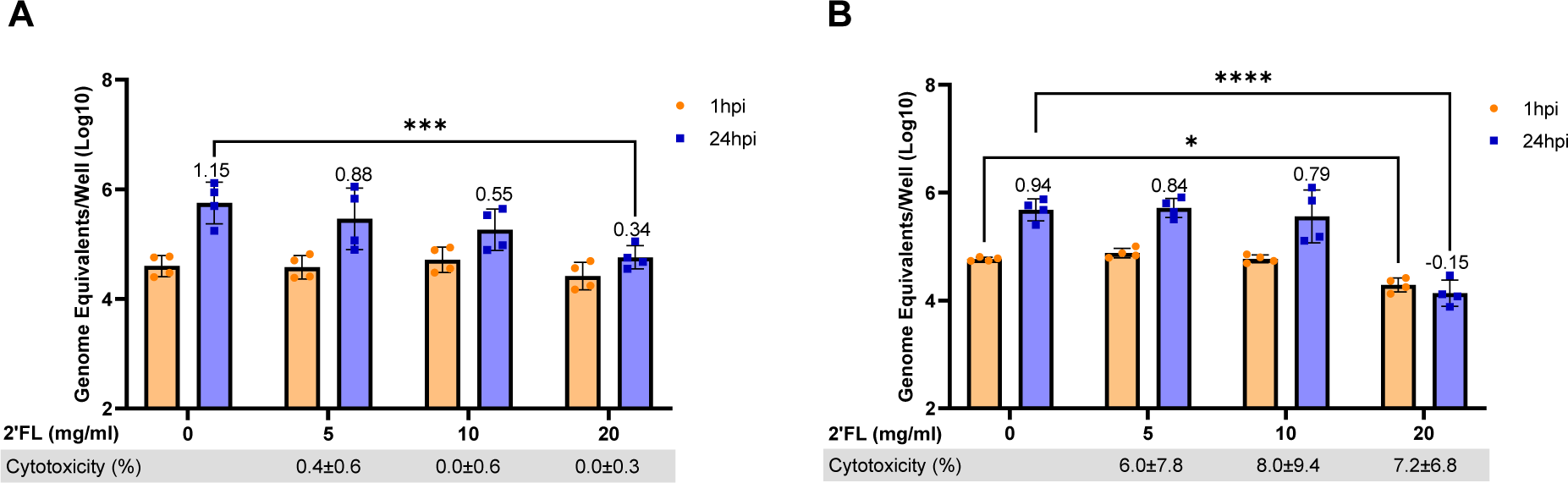
20 mg/ml 2’FL significantly reduces GII.4 Sydney [P16] HuNoV replication in adult duodenal HIE lines. Dose response assays were carried out in adult duodenal HIE lines (A) D109 and (B) D2004 using 5 mg/ml, 10 mg/ml and 20 mg/ml of 2’FL. GE/well were determined by RT-qPCR at 1 hour post infection (hpi) and 24 hpi. Numbers above the bars indicate log_10_ fold change comparing GE/well at 24 hpi to 1 hpi. Cytotoxicity (measured by lactase dehydrogenase assay) is represented in percentage below each graph. Data represented are means ± standard deviation (SD) from n=2 independent experiments with 2 technical replicates per experiment. The P-values were calculated using ANOVA, Sidak’s Multiple Comparisons Test. *P ≤ 0.05, ***P ≤ 0.001, ****P ≤ 0.0001.

### 2’FL SIGNIFICANTLY REDUCES GII.4 HUMAN NOROVIRUS REPLICATION IN ADULT JEJUNAL HIE LINES

To evaluate if the reduction in GII.4 Sydney [P16] replication with 2’FL could be seen in other intestinal segments, we next tested 2’FL in two adult jejunal HIE lines J2 and J11. Of note, we performed this and subsequent experiments only with 20 mg/ml of 2’FL since PGM-VLP assays and infectivity studies showed significant results only at the highest concentration. Both in J2 and J11 (Figure 3), GII.4 Sydney [P16] HuNoV showed ∼1.5log_10_ increase at 24 hpi compared to 1 hpi at baseline (0 mg/ml). 20 mg/ml of 2’FL showed a significant reduction in GII.4 Sydney [P16] HuNoV replication for both lines. When comparing HuNoV replication at 24 hpi, there was a 0.4log_10_ decrease in J2 and 0.7log_10_ decrease in J11. Similar to the duodenal HIEs, 20 mg/ml of 2’FL also showed a significant decrease in GII.4 Sydney [P16] binding for one jejunal HIE line (J2) but not the other. The 20 mg/ml of the HMO was not cytotoxic in either J2 or J11 HIEs.

**Figure 3:**
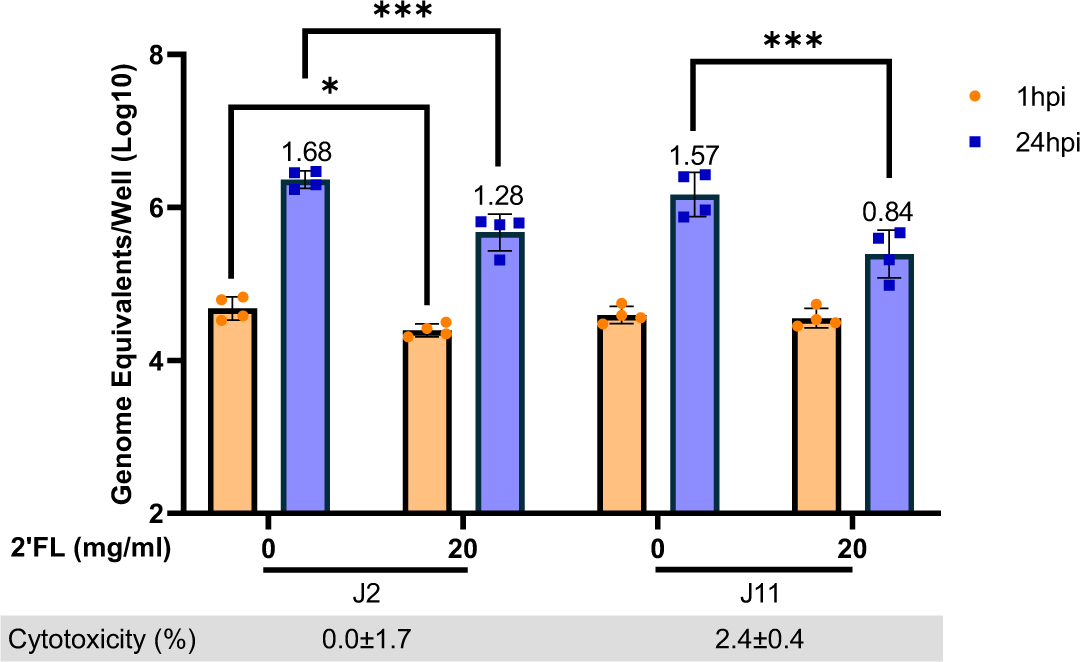
2’FL significantly reduces GII.4 Sydney [P16] HuNoV replication in adult jejunal HIE lines. 20 mg/ml of 2’FL was tested in two adult jejunal HIEs J2 and J11. GE/well were determined by RT-qPCR at 1 hpi and 24 hpi. Numbers above the bars indicate log_10_ fold change comparing GE/well at 24 hpi to 1 hpi. Cytotoxicity is represented in percentage below each graph. Data represented are means ± standard deviation (SD) from n=2 independent experiments with 2 technical replicates per experiment. The P-values were calculated using ANOVA, Sidak’s Multiple Comparisons Test. *P ≤ 0.05, ***P ≤ 0.001.

### 2’FL SIGNIFICANTLY REDUCES GII.4 HUMAN NOROVIRUS REPLICATION IN ALL INTESTINAL SEGMENTS OF THE SAME DONOR

The data shown above indicates 20 mg/ml 2’FL significantly inhibits GII.4 Sydney [P16] HuNoV replication in duodenal and jejunal HIEs. However, the magnitude of replication and inhibition varied between the different HIE lines. Since all the HIEs tested thus far were derived from different adult donors, it is possible that some of these differences could be attributed to variability between donors. We therefore wanted to evaluate the effect of 2’FL in intestinal segments from the same donor. 20 mg/ml of 2’FL was tested in a duodenal (D2004), jejunal (J2004) and ileal (I2004) segments from a single donor. Replication was highest in the ileum as measured by fold increases in GE/well from 1 hpi to 24 hpi, followed by jejunum and then duodenum (Figure 4). 20 mg/ml 2’FL significantly decreased both binding and replication of GII.4 Sydney [P16] HuNoV in all segments, with complete inhibition seen in the D2004 line. 2’FL was not cytotoxic in any of the segments.

**Figure 4:**
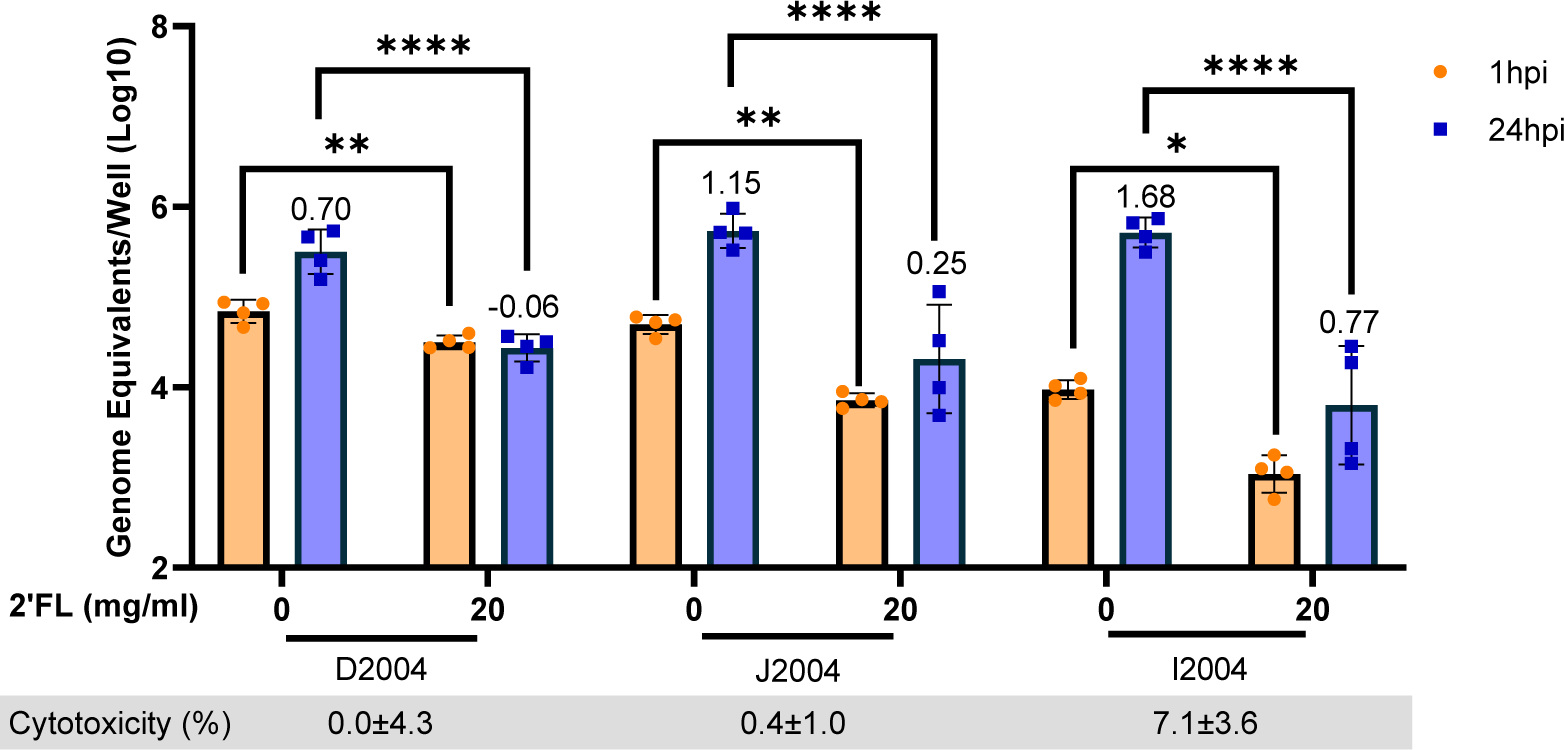
2’FL significantly reduces GII.4 Sydney [P16] HuNoV binding and replication in all segments from the same adult donor. Studies testing 20 mg/ml of 2’FL in duodenal (D2004), jejunal (J2004) and ileal (I2004) from an adult donor line. GE/well were determined by RT-qPCR at 1 hpi and 24 hpi. Numbers above the bars indicate log_10_ fold change comparing GE/well at 24 hpi to 1 hpi. Cytotoxicity is represented in percentage below each graph. Data represented are means ± standard deviation (SD) from n=2 independent experiments with 2 technical replicates per experiment. The P-values were calculated using ANOVA, Sidak’s Multiple Comparisons Test. *P ≤ 0.05, **P ≤ 0.01, ****P ≤ 0.0001.

### 2’FL SIGNIFICANTLY REDUCES GII.4 HUMAN NOROVIRUS REPLICATION IN PEDIATRIC DUODENAL BUT NOT INFANT JEJUNAL HIE LINES

As 2’FL significantly reduced HuNoV replication in adult lines, we next wanted to determine if similar outcomes would be observed in pediatric and infant HIE lines. Infectivity studies were carried out in two pediatric duodenal lines (4D and 8D, Figure 5A) and two infant jejunal lines (J1005 and J1006, Figure 5B). Similar to adult HIEs, higher replication in the absence of 2’FL was seen in infant jejunal HIEs (1.8log_10_) compared to pediatric duodenal HIEs (1.2log_10_). 20 mg/ml 2’FL reduced HuNoV replication, but not binding, in the two pediatric duodenal HIE lines. Surprisingly, when 20 mg/ml 2’FL was tested in two infant jejunal HIE lines (J1005 and J1006), no reduction of HuNoV binding or replication was observed (Figure 5B), suggesting that 2’FL is not acting as a decoy to block virus replication in these lines. 20 mg/ml 2’FL was not cytotoxic in both pediatric duodenal and infant jejunal lines.

**Figure 5:**
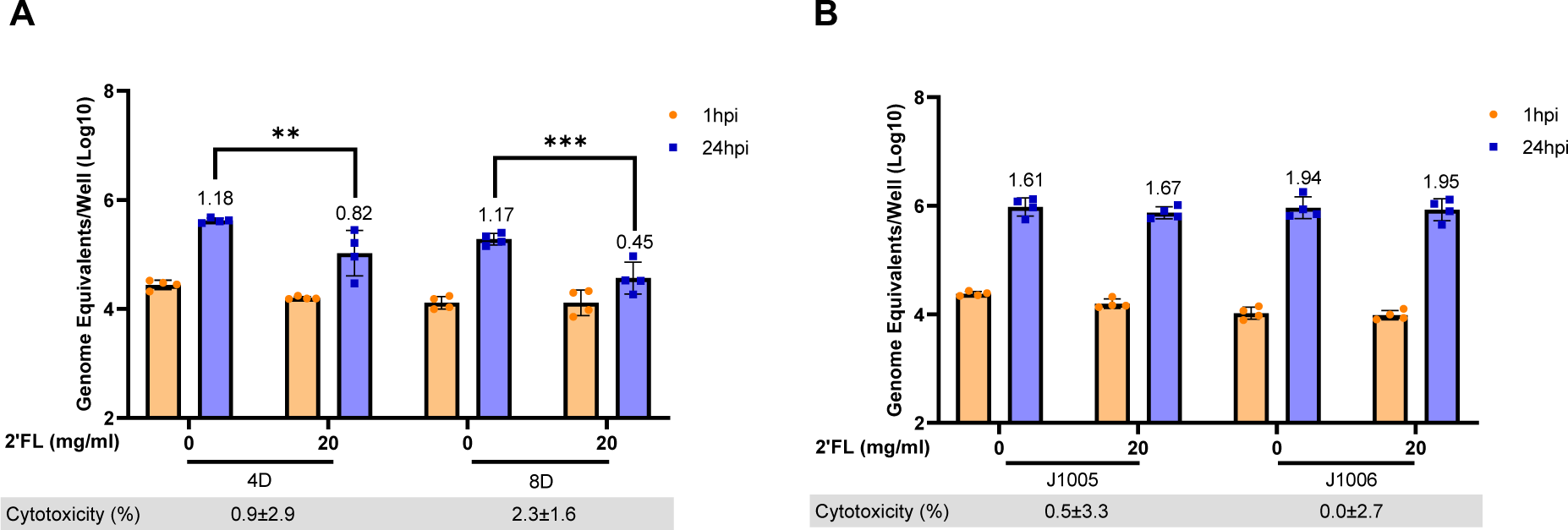
2’FL significantly reduces GII.4 Sydney [P16] HuNoV replication in pediatric duodenal but not infant jejunal HIE lines. 20 mg/ml of 2’FL was tested in (A) two pediatric duodenal HIEs (4D and 8D) and (B) two infant jejunal HIEs (J1005 and J1006). GEs per well were determined by RT-qPCR at 1 hpi and 24 hpi. Numbers above the bars indicate log_10_ fold change comparing GEs at 24 hpi to 1 hpi. Cytotoxicity is represented in percentage below each graph. Data represented are means ± standard deviation (SD) from n=2 independent experiments with 2 technical replicates per experiment. The P-values were calculated using ANOVA, Sidak’s Multiple Comparisons Test. **P ≤ 0.01, ***P ≤ 0.001.

### INFANT JEJUNAL LINES EXPRESS LOWER LEVEL OF α1-2-FUCOSYLATED HBGAS

As 20 mg/ml 2’FL didn’t inhibit GII.4 Sydney [P16] replication at 24 hpi in the infant jejunal lines but inhibited replication in the adult jejunal HIE lines tested, we wanted to evaluate if there was lower expression of fucosylated HBGAs in the infant lines. We compared the expression of α1-2-fucosylated glycans between the adult jejunal lines (J2 and J11) and infant jejunal lines (J1005 and J1006) by staining the HIEs with *Ulex europaeus* Agglutinin-1 (UEA-1, Figure 6A) (24). Significantly lower fluorescent intensity was observed in the infant jejunal lines compared to the adult jejunal lines (Figure 6B), suggesting the possibility of additional binding factors in the infant HIE lines other than α1-2-fucosylated HBGAs. There is no significant difference in fluorescent intensity between the two adult jejunal lines or the between the two infant jejunal lines (Figure 6B). The fluorescent intensities significantly correlate with levels of virus binding at 1hpi and with GE/TCID_50_ (Pearson r = 0.96 and 0.98, p<0.05, respectively).

**Figure 6:**
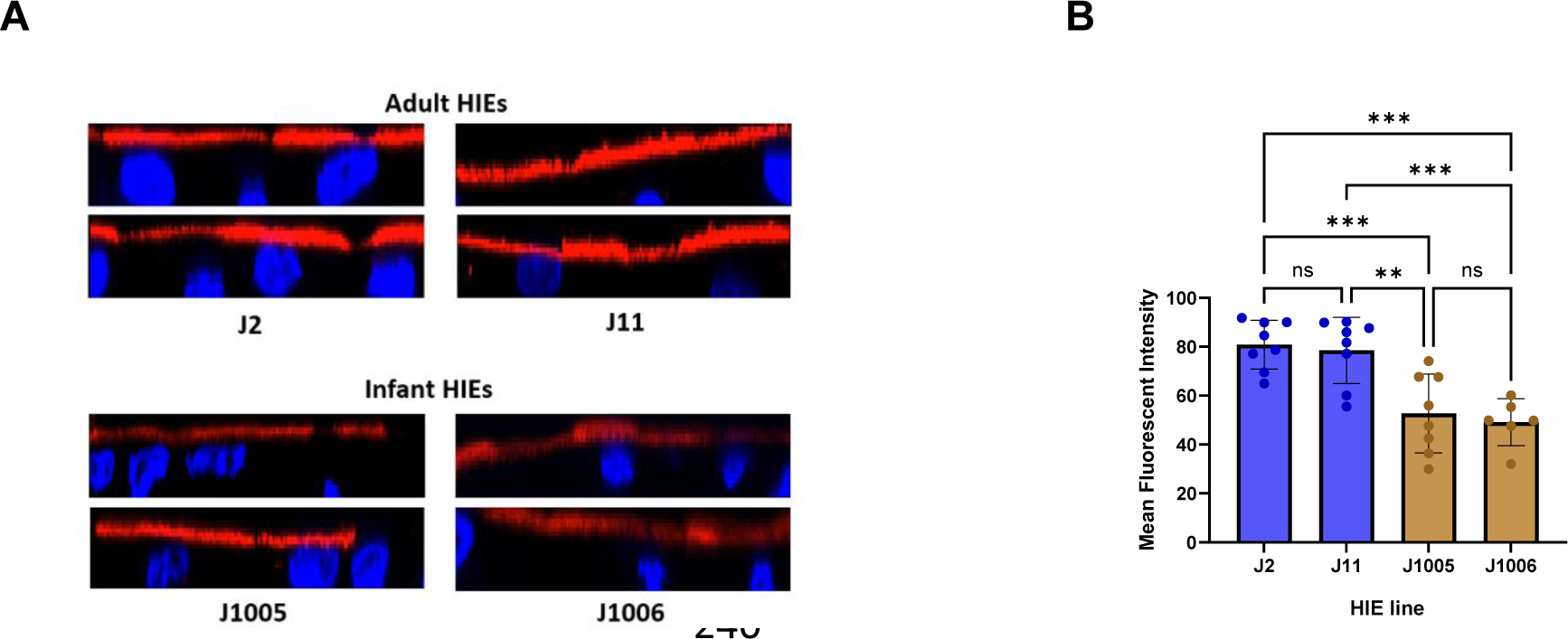
Level of HBGA expression is lower in infant jejunal HIE lines as compared to adult jejunal HIE lines. (A) Infant jejunal lines (J1005 and J1006) and adult jejunal lines (J2 and J11) stained with *Ulex europaeus* Agglutinin-1 (UEA-1) were imaged using confocal microscopy. Two representative images are shown per HIE line. (B) Fluorescence intensity was measured for each line using FIJI/Image J. Two-four fields per well were analyzed. Mean fluorescence data from 5 identical regions of interest (ROIs) per 2-4 fields were averaged. The P-values were calculated using ANOVA, Holm-Sidak’s Multiple Comparisons Test. **P ≤ 0.01, ***P ≤ 0.001. N=2 independent experiments.

## DISCUSSION

HMOs are known to act as decoy receptors for multiple enteric pathogens (25, 26). Previous studies have demonstrated that milk from secretor mothers, who produce α1-2-fucosylated HMOs, could block the binding of prototype Norwalk virus (GI.1) VLPs to intestinal tissues, H type I HBGA and saliva (27–29). Subsequent studies showed that 2’FL could block the binding of GI.1, GII.10 and GII.17 VLPs to PGM and saliva samples from multiple donors (19, 21). X-ray crystallography studies revealed that 2’FL binding occurred at the HBGA binding pockets suggesting that 2’FL can act as a decoy receptor for multiple HuNoV strains. However, data on 2’FL interactions with the globally dominant GII.4 genotype have been more variable. Two previous studies using VLPs from VA387 GII.4 strain suggested weak binding to 2’FL and the need for higher molecular weight glycoconjugates for inhibiting carbohydrate ligand interactions (30, 31). 2’FL at concentrations as high as 24 mg/ml did not inhibit GII.P16-GII.4 replication in zebrafish larvae although inhibition of binding to A-type saliva was seen (22). By contrast, structural studies suggest that the protruding domain of the GII.4 Sydney capsid protein binds 2’FL and HBGAs in the same pocket (20). In this study, we evaluated the effect of 2’FL on the infectivity of a recently circulating GII.4 Sydney [P16] HuNoV strain in HIEs. These nontransformed cultures serve as a physiologically relevant model system of the small intestinal epithelium and retain intestinal segment specificity as well as donor phenotypic characteristics. We discovered that 2’FL inhibits GII.4 Sydney [P16] HuNoV replication in multiple adult HIE lines without cytotoxicity (summarized in Table 2) and thus has the potential to be developed as a therapeutic for HuNoV gastroenteritis.

**Table 2:** Summary of the effect of 2’FL on GII.4 Sydney [P16] HuNoV replication. List of HIE lines, segment of origin and their respective age are shown. Average log_10_ fold increase in the absence of 2’FL and with 20 mg/ml 2’FL (percentage reduction) is shown.

| HIE | Segment | Age | Average fold increase<br>in the absence of<br>2'FL | Average fold<br>increase with<br>20 mg/ml 2'FL<br>(% reduction) |
| --- | --- | --- | --- | --- |
| D109 | Duodenal | 44 years | 1.15 | 0.34 (70.4%) |
| D2004 | Duodenal | 25 years | 0.82 | 0.105 (100%) |
| J2 | Jejunal | 52 years | 1.68 | 1.28 (23.8%) |
| J11 | Jejunal | 52 years | 1.57 | 0.84 (45.6%) |
| J2004 | Jejunal | 25 years | 1.15 | 0.25 (78.3%) |
| I2004 | Ileal | 25 years | 1.68 | 0.77 (54.2%) |
| 4D | Duodenal | 2 years | 1.18 | 0.82 (30.5%) |
| 8D | Duodenal | 5 years | 1.17 | 0.45 (61.5%) |
| J1005 | Jejunal | 10 weeks | 1.61 | 1.67 (0%) |
| J1006 | Jejunal | 12 weeks | 1.94 | 1.95 (0%) |

The concentration of 2’FL that inhibits GII.4 Sydney [P16] replication in HIEs is consistent with biochemical studies with GI.1, GII.17 and GII.10 VLPs where the IC_50_ was calculated to be between 5–20 g/L (21). While these concentrations are substantially higher than average concentrations in human milk, the safety profile of higher concentrations of 2’FL have been evaluated previously. A preclinical study in rats showed that oral administration of 2’FL up to 5000 mg per kilogram of body weight per day for over 90 days was not associated with any adverse effects based on clinical observations and histopathology, body weight gain and food consumption (32). A randomized, double-blind, placebo-controlled, oral supplementation study of 2’FL in 100 healthy adults showed that up to 20 g/day for about 12 days was safe and well tolerated (33). Microbiome composition analysis using 16S rRNA sequencing showed that HMO supplementation resulted in changes in the gut microbiota with increases in relative abundance of Actinobacteria and *Bifidobacterium*, and a reduction in relative abundance of Firmicutes and Proteobacteria. Chemical, chemo-enzymatic and enzymatic strategies to produce 2’FL have been described and include strategies for kilogram scale synthesis (34). Multiple 2’FL manufacturers, including the one for the 2’FL used in this study, have received “no questions” letters from the US Food and Drug Administration (FDA) regarding the generally recognized as safe (GRAS) notices for use of their HMO (35). Also, the European Food Safety Authority (EFSA) has published positive assessment opinions for use of 2’FL in food supplements. Infant formula supplemented with 2’FL is well-tolerated in healthy-term infants and supports age-appropriate growth (36–38). Additional health benefits of 2’FL have been described in various studies. Unbiased metabolomic analyses and short chain fatty acid production was evaluated in bioreactors seeded with fecal samples from 6 adults and 6 children (6 year old) that were supplemented with 0.5 – 1 g per day equivalent of 2’FL; these studies demonstrated significant increases in acetate and propionate production as well as aromatic lactic acids are linked to immune function (39). 2’FL was also associated with significant reduction in FITC-Dextran permeability in Caco2 cells and upregulation of tight junction proteins like Claudin-5 in colon-on-chip models under microfluidic conditions (40). Taken together, these data provide a promising outlook to regulatory pathways for clinical testing of 2’FL as an inhibitor for HuNoVs.

A critical observation in our study was that the inhibition of HuNoV replication varied by donor, intestinal segment, and age. The relative contribution of each of these factors remains to be elucidated. However, the availability of HIEs from all segments of the small intestine from a single donor allowed us to confirm that 2’FL can inhibit GII.4 Sydney [P16] replication across multiple segments. While the level of inhibition varied, with complete inhibition in duodenal HIEs to approximately 50% inhibition in ileal HIEs, it is to be noted that the magnitude of replication also varied, with the highest replication seen in ileal HIEs despite using 100 TCID_50_ of virus in all lines. The complete lack of inhibition in infant jejunal HIEs is particularly striking. We previously demonstrated significant transcriptional, morphological, and functional differences between the adult and infant jejunal HIEs used in this study (41). Of relevance to HMOs, the expression of lactase (β-galactosidase) was significantly higher in infant HIEs. However, previous studies have postulated that despite significant lactase presence, the upper small intestine of piglets and infants do not cleave HMOs (42). We evaluated differences in HBGA expression between infant and adult jejunal HIEs. The significantly lower expression of α1-2-fucosylated glycans on infant jejunal HIEs in comparison to adult lines suggests the possibility of additional cellular attachment factors on infant lines which allow viral infection and replication to occur despite the decoy activity of 2’FL.

Future studies can be performed to address some limitations of this work. First, additional mechanistic studies are required to determine if 2’FL only has decoy receptor activity or if host responses contribute to the antiviral effects. Second, while some lines show complete inhibition of GII.4 Sydney [P16] replication, the range of effects is large. Longer chain fucosylated HMOs like lacto-N-fucopentaose (LNFP) I or combinations of 2’FL with the other HMOs such as 3-fucosyllactose (3FL) can be tested as additional approaches to determine if consistent reduction in replication can be achieved across donors and segments. A recent structural study with nanobodies also demonstrated increased potency when used in combination with 2’FL (43). Such combination strategies could also be evaluated in future studies for effects on virus replication. Third, to evaluate the broad applicability of 2’FL or modified glycoconjugates, the effect on replication of additional HuNoV strains and in additional HIE lines in each age category/segment needs to be evaluated. Finally, assessment of 2’FL effects in HIEs from infants, toddlers and older children will allow us to determine whether there are developmentally regulated differences between receptor/co-receptor expression for HuNoVs.

We recently standardized a pipeline for evaluation of antivirals against HuNoVs using HIEs (23). We previously applied this pipeline to evaluate nitazoxanide, an anti-parasitic drug that is anecdotally used for the treatment of chronic HuNoV infections in immunocompromised patients. The present study demonstrates the utility of this pipeline to preclinically evaluate compounds based on known biology of HuNoVs. The study establishes the potential for 2’FL to be developed as a therapeutic for adults based on inhibition of virus replication. This is significant because previous studies have focused primarily on structural interactions and carbohydrate ligand blocking and did not demonstrate functional activity. Despite the high burden of disease, there are currently no approved antivirals or therapeutics for treating HuNoV infections, and a 2’FL-based therapeutic could have prophylactic applications in settings of high risk for outbreaks such as cruise ships or in treatment for acute or chronic infections.

## MATERIALS AND METHODS

### VLPS, VIRUS, AND 2’FL

GII.4 Sydney 2012 VLPs were used for the initial screening assay to evaluate whether 2’FL blocks the binding to PGM. VLPs were produced in a baculovirus system using open reading frame 2 (ORF2) + ORF3+ untranslated region (UTR) sequences (44). A GII.4 Sydney [P16] strain (isolate BCM 16-16, stock titer 4.26×106 GE/ul) was used for all infectivity experiments. 2’fucosyllactose, produced in bioengineered microbes, was generously provided in-kind by Jennewein GmbH, Germany, which was later acquired by Chr Hansen, Denmark, now part of Novonesis.

### HUMAN INTESTINAL ENTEROIDS

HIE lines from different intestinal segments and donors of different ages were used in this study. These include two adult duodenal lines (D109, D2004), three adult jejunal lines (J2, J11, J2004) and one adult ileal line (I2004). Of the adult lines, D2004, J2004 and I2004 were obtained from a single donor (45). In addition to HIE lines from adults, two pediatric duodenal lines (4D, 8D) and two infant jejunal (J1005, J1006) were included in this study. The ages of the HIE donors are listed in Table 1.

### HBGA BLOCKING ASSAYS

A 96-well polystyrene flat-bottom plate (Greiner Bio-One, 655001) was coated with 3 ug/ml PGM diluted in 0.01 M phosphate buffer saline (PBS) overnight at 4°C on a rocking platform. Following incubation, 1% non-fat dry milk (NFDM) in 100 mM sodium phosphate buffer (PB), pH 6.1 was added to the PGM-coated plate and incubated for 2 hours at room temperature protected from light. Meanwhile, two-fold dilutions of 2’FL ranging from 1.25 mg/ml to 20 mg/ml were incubated with 2.5 ug/ml GII.4 Sydney 2012 VLPs in a tissue-culture treated round bottom plate (Corning, 3799) at 4°C on a rocking platform for an hour. As the positive control, 2.5 ug/ml of GII.4 Sydney 2012 VLPs were diluted with PB buffer. Following incubation, the PGM coated plate was washed 5 times with cold PB buffer. The 2’FL-VLP solutions were transferred to the PGM coated plate and incubated at 4°C for 2 hours protected from light. Following incubation, the plate was washed 5 times with cold PB buffer. An in-house guinea pig anti-GII.4 Sydney primary antibody (1:3000) was added to the wells. The plate was incubated at 4°C for 1 hour protected from light. The plate was washed 5 times with cold PB buffer, and goat anti-guinea pig secondary antibody conjugated with HRP (1:5000, Sigma, A7289) was added to the wells. The plate was incubated at 4°C for 1 hour protected from light. After washing, TMB (3,3’,5,5’-Tetramethylbenzidine) substrate (KPL, 5120-0047), was added to all the wells for 10 minutes protected from light. 1M phosphoric acid was used as the stop solution and the absorbance was measured at 450nM using a microplate reader (Spectramax). The VLP binding assays were performed three times, with three technical replicate wells for each condition in an experiment.

### HUMAN NOROVIRUS INFECTIVITY

Three-dimensional (3D) HIE cultures were obtained from the Gastrointestinal Experimental Model Systems Core (GEMS) of the Texas Medical Center Digestive Diseases Center (TMC DDC) and plated as monolayers on 96-well plates as described previously using commercially available Intesticult™ Organoid Growth Medium (OGM) proliferation and differentiation media (46, 47). The GE/TCID_50_ was determined for each HIE line as described previously so that a standard dose of virus could be used across different HIE lines (23). 2’FL was diluted in OGM differentiation media with 500 μM sodium glycochenodeoxycholate (GCDCA; Sigma, G0759) and was added to 5-day differentiated HIE monolayers on a 96-well plate (Corning, 3595) with 100 TCID_50_ of virus. 100 TCID_50_ virus in the absence of 2’FL was used to determine baseline infectivity in the absence of treatment. Following incubation at 37°C for 1 hour, the HIE monolayers were washed 3 times with complete media without growth factors (CMGF-) and OGM differentiation media with 500 μM GCDCA was added to all the wells. The samples were incubated for a further 23 hours at 37°C. Total RNA was extracted using a KingFisher Flex machine (ThermoFisher) and MagMax-96 viral RNA isolated kit (Applied Biosystems) as described previously (47). RT-qPCR (Applied Biosystems) was carried out for the extracted RNA samples and viral replication was quantitated relative to a standard curve. GE/ul measured at 1 hpi and 24 hpi were used to estimate input virus and replication, respectively. The TCID_50_ assays were carried out twice for each line. Each infectivity experiment was performed twice with two technical replicate wells for each condition within an experiment. RT-qPCR assays were carried out using three technical replicates for each HIE well.

### CYTOTOXICITY ASSESSMENT

Cytotoxicity assays were carried out in tandem with the viral infectivity assays using the CytoTox 96® Non-Radioactive Cytotoxicity Assay (Promega, G1780). The assay was carried out according to the manufacturer’s instructions with some modifications wherein supernatants were diluted in media to achieve optical density (OD) values in the linear range of the assay (48). OD values were taken using a microplate reader at 490nM (Spectramax) and percent cytotoxicity was calculated for each sample.

### UEA-1 STAINING

5-day differentiated HIE monolayers plated on tissue culture treated slides (Ibidi, 80826) were fixed with 4% paraformaldehyde (Electron Microscopy Sciences, 15710-S) for 25 minutes at room temperature. The cells were incubated overnight at 4°C with Rhodamine-conjugated UEA-1 (Vector Laboratories, RL-1062-2) diluted 1:200 in 5% bovine serum albumin (BSA) in 0.01 M PBS + 0.1% triton (24). The cells were washed with 0.01 M PBS + 0.1% triton 3 times (10-minute incubations) and nuclei were stained with NucBlue Fixed Cell Stain ReadyProbes reagent (Invitrogen, R37606) diluted in 0.01 M PBS for 5 minutes. Orthogonal sections of the cells were imaged using a ZEISS confocal microscope (Laser Scanning Microscope LSM 980) using ZEISS ZEN 3.5 (blue edition) software. The images were further processed and analyzed using ImageJ2/FIJI. For quantifying fluorescence intensity, two to four fields per well were analyzed. Mean fluorescence data from 5 identical regions of interest (ROIs) per field were collected. The experiments were performed twice with two technical replicate wells in each experiment for each HIE line.

### STATISTICAL ANALYSIS

GraphPad Prism 9.5.1 was used for all statistical analyses. For the PGM-VLP assays, Student’s T-test was used to compare the 2’FL concentrations to the control. For the infectivity assays, comparison between 1 hpi and 24 hpi groups in the presence and absence of 2’FL was performed using a two-way ANOVA and Sidak’s post-hoc multiple comparisons analyses. For comparing fluorescent intensities of UEA-1 staining in the immunofluorescence assays between the different lines, a one-way ANOVA was performed using Holm-Sidak’s multiple comparisons test for post-hoc analyses. Error bars denote standard deviation (SD) for all graphs.

## ACKNOWLEDGMENTS

We thank Dr. Mark Donowitz (Johns Hopkins University Medical School) for providing the two pediatric duodenal HIE lines. We thank Xei-Li Zeng, Yi-Ting Shen, and Aaya Boussattach from the Texas Medical Center Digestive Diseases Center (TMC DDC) Gastrointestinal Experimental Model Systems (GEMS) core (supported by the NIH P30 DK056338 grant) for assistance with the maintenance and plating of human intestinal enteroids. This work was supported by a Pilot/Feasibility grant from TMC DDC (SR) and by the Public Health Service grant P01 AI 057788 (M.K.E and R.L.A.). The purchase of the Zeiss Laser Scanning Microscope LSM 980 with Airyscan 2 used for microscopy studies was supported by the S10 OD028480 grant.

## REFERENCES

1. Ahmed SM, Hall AJ, Robinson AE, Verhoef L, Premkumar P, Parashar UD, Koopmans M, Lopman BA. 2014. Global prevalence of norovirus in cases of gastroenteritis: a systematic review and meta-analysis. Lancet Infect Dis 14:725–730.

2. Pires SM, Fischer-Walker CL, Lanata CF, Devleesschauwer B, Hall AJ, Kirk MD, Duarte AS, Black RE, Angulo FJ. 2015. Aetiology-Specific Estimates of the Global and Regional Incidence and Mortality of Diarrhoeal Diseases Commonly Transmitted through Food. PLoS One 10:e0142927.

3. Lopman BA, Steele D, Kirkwood CD, Parashar UD. 2016. The Vast and Varied Global Burden of Norovirus: Prospects for Prevention and Control. PLoS Med 13:e1001999.

4. Gaythorpe KAM, Trotter CL, Lopman B, Steele M, Conlan AJK. 2018. Norovirus transmission dynamics: a modelling review. Epidemiol Infect 146:147–158.

5. Bartsch SM, Lopman BA, Ozawa S, Hall AJ, Lee BY. 2016. Global Economic Burden of Norovirus Gastroenteritis. PLoS One 11:e0151219.

6. Atmar RL, Ramani S, Estes MK. 2018. Human noroviruses: recent advances in a 50-year history. Curr Opin Infect Dis 31:422–432.

7. Huang P, Farkas T, Zhong W, Tan M, Thornton S, Morrow AL, Jiang X. 2005. Norovirus and histo-blood group antigens: demonstration of a wide spectrum of strain specificities and classification of two major binding groups among multiple binding patterns. J Virol 79:6714–22.

8. Taube S, Mallagaray A, Peters T. 2018. Norovirus, glycans and attachment. Curr Opin Virol 31:33–42.

9. Cheng YJ, Yeung CY. 2021. Recent advance in infant nutrition: Human milk oligosaccharides. Pediatr Neonatol 62:347–353.

10. Berger PK, Ong ML, Bode L, Belfort MB. 2023. Human Milk Oligosaccharides and Infant Neurodevelopment: A Narrative Review. Nutrients 15.

11. Plows JF, Berger PK, Jones RB, Alderete TL, Yonemitsu C, Najera JA, Khwajazada S, Bode L, Goran MI. 2021. Longitudinal Changes in Human Milk Oligosaccharides (HMOs) Over the Course of 24 Months of Lactation. J Nutr 151:876–882.

12. Bode L. 2012. Human milk oligosaccharides: every baby needs a sugar mama. Glycobiology 22:1147–62.

13. Yu ZT, Nanthakumar NN, Newburg DS. 2016. The Human Milk Oligosaccharide 2’-Fucosyllactose Quenches Campylobacter jejuni-Induced Inflammation in Human Epithelial Cells HEp-2 and HT-29 and in Mouse Intestinal Mucosa. J Nutr 146:1980–1990.

14. Wiese M, Schuren FHJ, Smits WK, Kuijper EJ, Ouwens A, Heerikhuisen M, Vigsnaes L, van den Broek TJ, de Boer P, Montijn RC, van der Vossen J. 2022. 2’-Fucosyllactose inhibits proliferation of Clostridioides difficile ATCC 43599 in the CDi-screen, an in vitro model simulating Clostridioides difficile infection. Front Cell Infect Microbiol 12:991150.

15. Wang Y, Zou Y, Wang J, Ma H, Zhang B, Wang S. 2020. The Protective Effects of 2’-Fucosyllactose against E. Coli O157 Infection Are Mediated by the Regulation of Gut Microbiota and the Inhibition of Pathogen Adhesion. Nutrients 12.

16. Laucirica DR, Triantis V, Schoemaker R, Estes MK, Ramani S. 2017. Milk Oligosaccharides Inhibit Human Rotavirus Infectivity in MA104 Cells. J Nutr 147:1709–1714.

17. Lou F, Hu R, Chen Y, Li M, An X, Song L, Tong Y, Fan H. 2022. 2’-Fucosyllactose Inhibits Coxsackievirus Class A Type 9 Infection by Blocking Virus Attachment and Internalisation. Int J Mol Sci 23.

18. Yu W, Li Y, Liu D, Wang Y, Li J, Du Y, Gao GF, Li Z, Xu Y, Wei J. 2023. Evaluation and Mechanistic Investigation of Human Milk Oligosaccharide against SARS-CoV-2. J Agric Food Chem 71:16102–16113.

19. Weichert S, Koromyslova A, Singh BK, Hansman S, Jennewein S, Schroten H, Hansman GS. 2016. Structural Basis for Norovirus Inhibition by Human Milk Oligosaccharides. J Virol 90:4843–4848.

20. Schroten H, Hanisch FG, Hansman GS. 2016. Human Norovirus Interactions with Histo-Blood Group Antigens and Human Milk Oligosaccharides. J Virol 90:5855–5859.

21. Koromyslova A, Tripathi S, Morozov V, Schroten H, Hansman GS. 2017. Human norovirus inhibition by a human milk oligosaccharide. Virology 508:81–89.

22. Tan MTH, Li Y, Eshaghi Gorji M, Gong Z, Li D. 2021. Fucoidan But Not 2’-Fucosyllactose Inhibits Human Norovirus Replication in Zebrafish Larvae. Viruses 13.

23. Lewis MA, Cortes-Penfield NW, Ettayebi K, Patil K, Kaur G, Neill FH, Atmar RL, Ramani S, Estes MK. 2023. Standardization of an antiviral pipeline for human norovirus in human intestinal enteroids demonstrates nitazoxanide has no to weak antiviral activity. Antimicrob Agents Chemother 67:e0063623.

24. Haga K, Ettayebi K, Tenge VR, Karandikar UC, Lewis MA, Lin SC, Neill FH, Ayyar BV, Zeng XL, Larson G, Ramani S, Atmar RL, Estes MK. 2020. Genetic Manipulation of Human Intestinal Enteroids Demonstrates the Necessity of a Functional Fucosyltransferase 2 Gene for Secretor-Dependent Human Norovirus Infection. mBio 11.

25. Moore RE, Xu LL, Townsend SD. 2021. Prospecting Human Milk Oligosaccharides as a Defense Against Viral Infections. ACS Infect Dis 7:254–263.

26. Triantis V, Bode L, van Neerven RJJ. 2018. Immunological Effects of Human Milk Oligosaccharides. Front Pediatr 6:190.

27. Ruvoen-Clouet N, Mas E, Marionneau S, Guillon P, Lombardo D, Le Pendu J. 2006. Bile-salt-stimulated lipase and mucins from milk of ‘secretor’ mothers inhibit the binding of Norwalk virus capsids to their carbohydrate ligands. Biochem J 393:627–34.

28. Jiang X, Huang P, Zhong W, Tan M, Farkas T, Morrow AL, Newburg DS, Ruiz-Palacios GM, Pickering LK. 2004. Human milk contains elements that block binding of noroviruses to human histo-blood group antigens in saliva. J Infect Dis 190:1850–9.

29. Marionneau S, Ruvoen N, Le Moullac-Vaidye B, Clement M, Cailleau-Thomas A, Ruiz-Palacois G, Huang P, Jiang X, Le Pendu J. 2002. Norwalk virus binds to histo-blood group antigens present on gastroduodenal epithelial cells of secretor individuals. Gastroenterology 122:1967–77.

30. Huang P, Morrow AL, Jiang X. 2009. The carbohydrate moiety and high molecular weight carrier of histo-blood group antigens are both required for norovirus-receptor recognition. Glycoconj J 26:1085–96.

31. Shang J, Piskarev VE, Xia M, Huang P, Jiang X, Likhosherstov LM, Novikova OS, Newburg DS, Ratner DM. 2013. Identifying human milk glycans that inhibit norovirus binding using surface plasmon resonance. Glycobiology 23:1491–8.

32. Coulet M, Phothirath P, Allais L, Schilter B. 2014. Pre-clinical safety evaluation of the synthetic human milk, nature-identical, oligosaccharide 2’-O-Fucosyllactose (2’FL). Regul Toxicol Pharmacol 68:59–69.

33. Elison E, Vigsnaes LK, Rindom Krogsgaard L, Rasmussen J, Sorensen N, McConnell B, Hennet T, Sommer MO, Bytzer P. 2016. Oral supplementation of healthy adults with 2’-O-fucosyllactose and lacto-N-neotetraose is well tolerated and shifts the intestinal microbiota. Br J Nutr 116:1356–1368.

34. Agoston K, Hederos MJ, Bajza I, Dekany G. 2019. Kilogram scale chemical synthesis of 2’-fucosyllactose. Carbohydr Res 476:71–77.

35. Zhu Y, Wan L, Li W, Ni D, Zhang W, Yan X, Mu W. 2022. Recent advances on 2’-fucosyllactose: physiological properties, applications, and production approaches. Crit Rev Food Sci Nutr 62:2083–2092.

36. Marriage BJ, Buck RH, Goehring KC, Oliver JS, Williams JA. 2015. Infants Fed a Lower Calorie Formula With 2’FL Show Growth and 2’FL Uptake Like Breast-Fed Infants. J Pediatr Gastroenterol Nutr 61:649–58.

37. Lasekan J, Choe Y, Dvoretskiy S, Devitt A, Zhang S, Mackey A, Wulf K, Buck R, Steele C, Johnson M, Baggs G. 2022. Growth and Gastrointestinal Tolerance in Healthy Term Infants Fed Milk-Based Infant Formula Supplemented with Five Human Milk Oligosaccharides (HMOs): A Randomized Multicenter Trial. Nutrients 14.

38. Puccio G, Alliet P, Cajozzo C, Janssens E, Corsello G, Sprenger N, Wernimont S, Egli D, Gosoniu L, Steenhout P. 2017. Effects of Infant Formula With Human Milk Oligosaccharides on Growth and Morbidity: A Randomized Multicenter Trial. J Pediatr Gastroenterol Nutr 64:624–631.

39. Bajic D, Wiens F, Wintergerst E, Deyaert S, Baudot A, Van den Abbeele P. 2023. HMOs Exert Marked Bifidogenic Effects on Children’s Gut Microbiota Ex Vivo, Due to Age-Related Bifidobacterium Species Composition. Nutrients 15.

40. Suligoj T, Vigsnaes LK, Abbeele PVD, Apostolou A, Karalis K, Savva GM, McConnell B, Juge N. 2020. Effects of Human Milk Oligosaccharides on the Adult Gut Microbiota and Barrier Function. Nutrients 12.

41. Adeniyi-Ipadeola GO, Hankins JD, Kambal A, Zeng XL, Patil K, Poplaski V, Bomidi C, Nguyen-Phuc H, Grimm SL, Coarfa C, Stossi F, Crawford SE, Blutt SE, Speer AL, Estes MK, Ramani S. 2024. Infant and Adult Human Intestinal Enteroids are Morphologically and Functionally Distinct. In press.

42. Engfer MB, Stahl B, Finke B, Sawatzki G, Daniel H. 2000. Human milk oligosaccharides are resistant to enzymatic hydrolysis in the upper gastrointestinal tract. Am J Clin Nutr 71:1589–96.

43. Ruoff K, Kilic T, Devant J, Koromyslova A, Ringel A, Hempelmann A, Geiss C, Graf J, Haas M, Roggenbach I, Hansman G. 2019. Structural Basis of Nanobodies Targeting the Prototype Norovirus. J Virol 93.

44. Ayyar BV, Ettayebi K, Salmen W, Karandikar UC, Neill FH, Tenge VR, Crawford SE, Bieberich E, Prasad BVV, Atmar RL, Estes MK. 2023. CLIC and membrane wound repair pathways enable pandemic norovirus entry and infection. Nat Commun 14:1148.

45. Ettayebi K, Kaur G, Patil K, Dave J, Ayyar BV, Tenge VR, Neill FH, Zeng XL, Speer AL, Dirienzi S, Britton RA, Blutt SE, Crawford SE, Ramani S, Atmar RL, Estes MK. Advancements in Human Norovirus Cultivation in Human Intestinal Enteroids. bioRxiv 2024.05.24.595764.

46. Tenge V, Vijayalakshmi Ayyar B, Ettayebi K, Crawford SE, Shen YT, Neill FH, Atmar RL, Estes MK. 2024. Bile acid-sensitive human norovirus strains are susceptible to sphingosine-1-phosphate receptor 2 inhibition. In press.

47. Ettayebi K, Tenge VR, Cortes-Penfield NW, Crawford SE, Neill FH, Zeng XL, Yu X, Ayyar BV, Burrin D, Ramani S, Atmar RL, Estes MK. 2021. New Insights and Enhanced Human Norovirus Cultivation in Human Intestinal Enteroids. mSphere 6.

48. Lewis MA PK, Ettayebi K, Estes MK, Atmar RL, Ramani S. Divergent responses of human intestinal organoid monolayers using commercial in vitro cytotoxicity assays. In press.

